# Evolution of DNA methylation in *Papio* baboons

**DOI:** 10.1101/400093

**Authors:** Tauras Vilgalys, Jeffrey Rogers, Clifford Jolly, Baboon Genome Analysis Consortium, Sayan Mukherjee, Jenny Tung

**Affiliations:** Department of Evolutionary Anthropology, Duke University, Durham, NC; Human Genome Sequencing Center, Baylor College of Medicine, Houston, TX; Department of Molecular and Human Genetics, Baylor College of Medicine, Houston, TX; Department of Anthropology, New York University, New York, NY; Center for the Study of Human Origins, New York University, New York, NY; New York Consortium for Evolutionary Primatology; Department of Statistical Science, Duke University, Durham, NC; Department of Mathematics, Duke University, Durham, NC; Department of Computer Science, Duke University, Durham, NC; Department of Biology, Duke University, Durham, NC; Duke University Population Research Institute, Duke University, Durham, NC; Institute of Primate Research, National Museums of Kenya, Karen, Nairobi, Kenya

## Abstract

Changes in gene regulation have long been thought to play an important role in primate evolution. However, although a number of studies have compared genome-wide gene expression patterns across primate species, fewer have investigated the gene regulatory mechanisms that underlie such patterns, or the relative contribution of drift versus selection. Here, we profiled genome-scale DNA methylation levels from five of the six extant species of the baboon genus *Papio* (4–14 individuals per species). This radiation presents the opportunity to investigate DNA methylation divergence at both shallow and deeper time scales (380,000 – 1.4 million years). In contrast to studies in human populations, but similar to studies in great apes, DNA methylation profiles clearly mirror genetic and geographic structure. Divergence in DNA methylation proceeds fastest in unannotated regions of the genome and slowest in regions of the genome that are likely more constrained at the sequence level (e.g., gene exons). Both heuristic approaches and Ornstein-Uhlenbeck models suggest that DNA methylation levels at a small set of sites have been affected by positive selection, and that this class is enriched in functionally relevant contexts, including promoters, enhancers, and CpG islands. Our results thus indicate that the rate and distribution of DNA methylation changes across the genome largely mirror genetic structure. However, at some CpG sites, DNA methylation levels themselves may have been a target of positive selection, pointing to loci that could be important in connecting sequence variation to fitness-related traits.

## Introduction

Changes in gene regulation have long been hypothesized to play an important role in trait evolution (Britten and Davidson 1971; King and Wilson 1975; Jacob 1977; Wray 2007; Stern and Orgogozo 2008). Regulatory changes have the potential to be more modular, and hence more specific to the individual tissues, environmental conditions, or developmental time points targeted by selection, than protein-coding changes (Stern 2000). In addition, regulatory regions are believed to have larger mutational target sizes, increasing the rate at which they may evolve (Landry, et al. 2007). In support of the importance of regulatory evolution, a number of studies have identified regulatory changes that contribute to species-specific adaptations. For example, non-coding variants that regulate the *ectodysplasin* and *pitx1* genes underlie morphological changes that separate saltwater threespine sticklebacks (*Gasterosteus aculeatus)* from their close freshwater relatives (Colosimo, et al. 2004; Shapiro, et al. 2004; Colosimo, et al. 2005). Similarly, wing pattern mimicry in *Heliconius* butterflies has been repeatedly shaped by regulatory evolution near the *optix* gene, in which convergent changes at different *cis*-regulatory variants have produced similar patterns of wing coloration (Reed, et al. 2011; Heliconius Genome Consortium 2012). Together, these and other case studies (e.g. Abzhanov, et al. 2004; Prud’Homme, et al. 2006; Manceau, et al. 2011; Jones, et al. 2012; Poelstra, et al. 2014) provide compelling examples of the importance of regulatory sequence changes to adaptive evolution.

However, evaluating the role of gene regulation in adaptive trait evolution also requires understanding the genome-wide distribution of selectively relevant regulatory variants. To address this question, two approaches have commonly been employed: sequence-based tests for selection and comparative analyses of gene expression phenotypes themselves. The first approach has identified signatures of natural selection in regulatory regions both within and between species (e.g., Pollard, et al. 2006; Prabhakar, et al. 2006; Kosiol, et al. 2008). In primates, for example, genes associated with developmental or neuronal functions have been argued to contain more signatures of positive selection in noncoding regions than in their coding sequences (Haygood, et al. 2010). Relative to other genetic variants, loci that affect gene expression in humans also have larger integrated haplotype scores, providing evidence for recent positive selection (Nédélec, et al. 2016; Kim-Hellmuth, et al. 2017). Consistent with these findings, variants associated with disease risk, fecundity, and other selectively relevant traits are often found within non-coding regions, and likely affect gene expression levels (Nicolae, et al. 2010; Wray 2013).

The second approach investigates patterns of gene expression across species to search for cases consistent with adaptive evolution. Several patterns have emerged from this work. First, overall differences in gene expression accumulate over evolutionary time, such that more closely related species have more similar gene expression profiles. Global clustering approaches from the same tissue thus tend to faithfully reproduce the species phylogeny (Brawand, et al. 2011; Sudmant, et al. 2015), and exceptions to this pattern suggest possible cases of natural selection. For example, gene expression levels in testis, but not in other tissues, group humans and gorillas to the exclusion of chimpanzees and bonobos (Brawand, et al. 2011). This pattern is consistent with elevated sexual selection on male reproductive physiology in chimpanzees and bonobos, which are characterized by unusually large testis to body size ratios relative to other primates (Schultz 1938). Second, stabilizing selection appears to constrain most gene expression levels. Comparative analyses of gene expression have found that most genes are characterized by low levels of intra- and inter-specific divergence, a pattern consistent with stabilizing selection (Rifkin, et al. 2003; Gilad, et al. 2006a; Khaitovich, et al. 2006; Blekhman, et al. 2008; Coolon et al. 2014; Hodgins-Davis et al. 2015). Furthermore, within species, regulatory variants of large effect tend to have low allele frequencies, suggesting that they are typically selected against (Battle, et al. 2014; Hernandez, et al. 2017; Schoech, et al. 2017). In support of this argument, experimental mutation accumulation lines exhibit an excess of gene expression variation compared to that observed in natural populations. They also accumulate differences in gene expression at a faster rate than observed in between-species comparisons (Denver, et al. 2005; Rifkin, et al. 2005).

Thus, both sequence-based studies and comparative studies of gene expression support a central role for selection on gene expression evolution, dominated by stabilizing selection but with an additional contribution made by positive selection (Signor and Nuzhdin 2018). However, gene expression patterns themselves are a product of multiple underlying regulatory mechanisms, which govern chromatin accessibility, transcription factor binding, and mRNA processing, splicing, and stability. These mechanisms link genetic variation in DNA sequence to selectively relevant gene expression phenotypes (Gallego Romero, et al. 2012; Pai and Gilad 2014). For example, in humans, genetic variants that affect chromatin accessibility and DNA methylation often affect gene expression as well, indicating that these mechanisms functionally link DNA sequence variation to gene expression (Degner, et al. 2012; Banovich, et al. 2014; Gate, et al. 2018). Between species, however, we know considerably less about how gene regulatory mechanisms evolve, including their relative contributions to lineage-specific shifts in gene expression levels (Pai and Gilad 2014).

Comparative studies to date have focused most intensively on DNA methylation, an epigenetic regulatory mechanism that refers to the covalent addition of a methyl group to a cytosine base and that can affect transcription factor binding, chromatin accessibility, and gene expression (Klose and Bird 2006; Weber, et al. 2007; Jones 2012; but see also Shibata, et al. 2012; Zhou, et al. 2014; Villar, et al. 2015; Berthelot, et al. 2018 for work on other mechanisms). In primates, comparisons between humans, chimpanzees, and rhesus macaques suggest that divergence in DNA methylation is associated with changes in gene expression (Zeng, et al. 2012; Heyn, et al. 2013), explaining 15-21% of expression differences between species (Pai, et al. 2011). Like gene expression, divergence in DNA methylation also increases with genetic distance (Hernando-Herraez, et al. 2013). However, comparisons among human populations suggest that DNA methylation evolves in a more clock-like fashion than gene expression, possibly because gene expression phenotypes evolve under greater functional constraint (Carja, et al. 2017). Unlike for gene expression levels (Rifkin, et al. 2003; Gilad, et al. 2006a; Khaitovich, et al. 2006; Whitehead and Crawford 2006; Blekhman, et al. 2010; Brawand, et al. 2011; Rohlfs and Nielsen 2014), the relative contribution of genetic drift and natural selection to DNA methylation evolution across species has not been investigated.

Here, we address this gap by investigating the evolution of genome-wide DNA methylation levels in the baboon genus *Papio*. Baboons radiated in sub-Saharan Africa over the past 1.4 million years to include six currently recognized extant species: anubis baboons (*P. anubis*, also called the olive baboon), hamadryas baboons (*P. hamadryas*), and Guinea baboons (*P. papio*) in the northern half of Africa and the Arabian peninsula; and yellow baboons (*P. cynocephalus*), chacma baboons (*P. ursinus*), and Kinda baboons (*P. kindae*) in central and southern Africa (Fig. 1A: Jolly 1993; Rogers, et al. *in review*). Studying DNA methylation divergence in this species complex thus provides additional resolution on the rate of DNA methylation evolution in primates, as previous studies have concentrated either on deeply diverged great apes (5 – 15 million years of divergence) or on closely related human populations (Pai, et al. 2011; Hernando-Herraez, et al. 2013; Heyn, et al. 2013; Hernando-Herraez, et al. 2015; Mendizabal, et al. 2016; Carja, et al. 2017). Genetic evidence indicates that branching events leading to the extant baboon species occurred on an intermediate time-scale, between 0.380 and 1.4 – 2.0 million years ago (Zinner, et al. 2013; Rogers, et al. *in review*). Further, because baboon genetic diversity is unusually well-characterized (Wall, et al. 2016; Leffler 2017), focusing on baboons also allowed us to investigate the relationship between DNA methylation and patterns of genetic variation across the genome.

**Figure 1.**
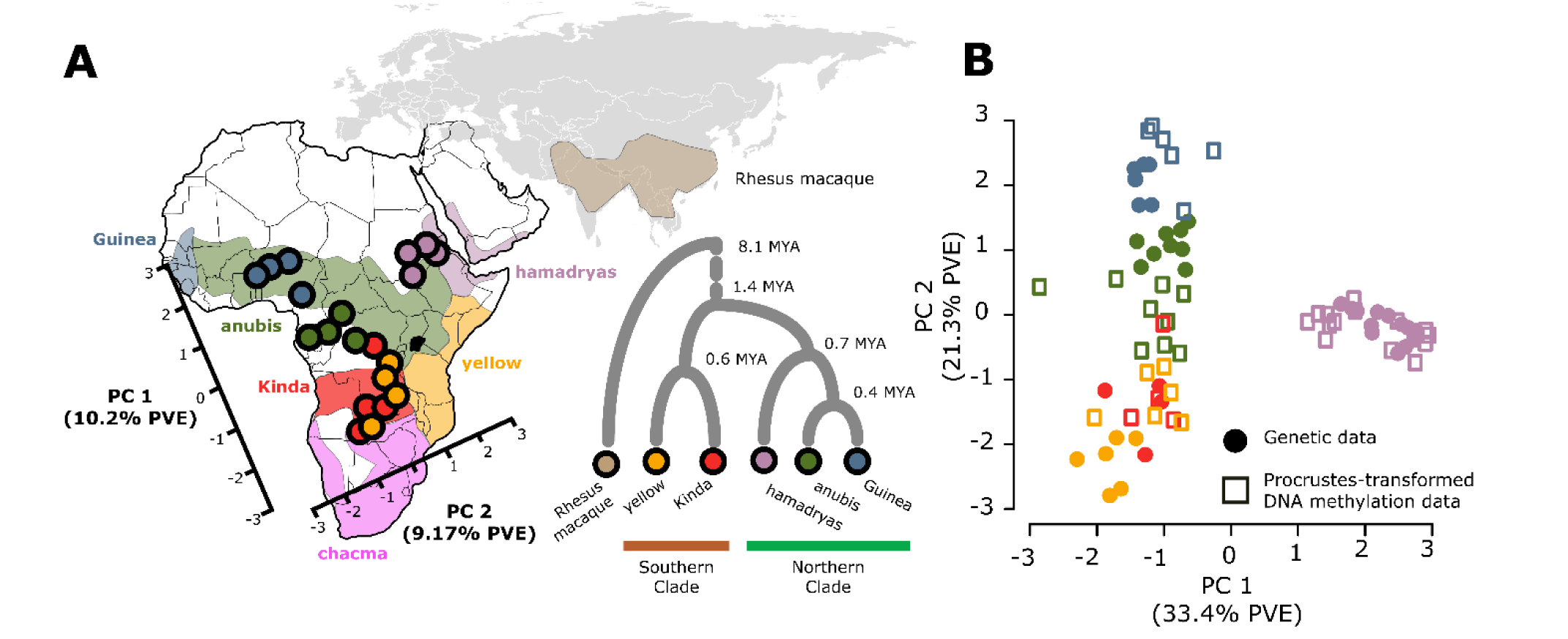
Geographic and genetic structure in baboon DNA methylation patterns. **(A)** The first two principal components from a PCA of baboon DNA methylation profiles (subsampled to n=4 individuals per species) projected onto the geographic distribution of baboon species in Africa. Northern clade (cool colors) and southern clade (warm colors) baboons separate along the first PC. Distribution of the six commonly recognized baboon allotaxa in Africa and the Arabian peninsula is based on Zinner, et al. (2013) and modified from a map created by Kenneth Chiou (CC BY 3.0 license); note that points reflect coordinates for DNA methylation data in PC space, not sampling location. Phylogenetic relationships between the five species included in this data set, with rhesus macaque as an outgroup, are shown in the inset (divergence dates within baboons from Rogers, et al. (*in review*) and between baboons and macaques from Perelman et al. (2011). **(B)** Procrustes transformation of PCs 1 and 2 of the DNA methylation data (empty squares) conforms with PCs 1 and 2 of genotype data (solid circles) from the same samples (Procrustes *t*_0_ = 0.89, p < 10^−6^). PVE values on the x and y axis are provided for the genotype data.

To do so, we generated genome-scale bisulfite sequencing data for 4 – 14 members of each of five of the extant species (all but chacma baboons). We asked: (i) to what degree does phylogenetic divergence between baboon species predict evolutionary change in DNA methylation levels? (ii) how are clade- and species-specific shifts in DNA methylation distributed across the baboon genome? and (iii) what are the relative contributions of natural selection and genetic drift to patterns of DNA methylation across species? Our results show that divergence in DNA methylation is closely linked to genetic divergence in baboons. Additionally, heterogeneity in DNA methylation divergence is explained by a combination of functional context, mean methylation level, and differences in selective constraint. At a subset of sites, these differences are consistent with lineage-specific selective shifts, suggesting candidate loci for which interspecific changes in gene expression may be explained by selection on DNA methylation.

## Results

### Genome-wide variation in DNA methylation reflects geography and phylogenetic structure

We generated DNA methylation profiles for 39 baboons and 5 rhesus macaques (*Macaca mulatta*) (Table S1) using reduced representation bisulfite sequencing (RRBS: Gu, et al. 2011; Boyle, et al. 2012). After filtering for CpG sites where at least half of our study subjects were sequenced at a mean coverage of at least 5x, the data set included DNA methylation estimates for 2,450,153 CpG sites throughout the genome. As expected for RRBS data, these sites were strongly enriched in or near CpG dense regions of the genome, including CpG islands, CpG shores, gene bodies, and promoters (Fig. S1). At least one CpG site in the promoter or gene body was included for 75.2% of Ensembl-annotated protein-coding genes in the reference anubis baboon genome (*Panu2.0*; Fig. S1). To investigate patterns of DNA methylation variation across *Papio,* we subsequently focused on the subset of 756,262 CpG sites that were not constitutively hyper- or hypo-methylated (mean methylation level ∈ [10%,90%]across all study subjects).

Two of the species we sampled (hamadryas baboons and anubis baboons) included individuals from multiple source populations (Table S1). However, because source population was not significantly associated with variation in DNA methylation within species (Supplementary Methods), we grouped all samples from the same species together for subsequent analysis.

To investigate the relationship between DNA methylation levels and genetic divergence, we first performed principal components analysis (PCA) on the DNA methylation data. With rhesus macaques included, the first principal component explained 14% of the overall variance in the data and separated all baboons from all rhesus macaques (Fig. S2). Subsequent PCs captured variation within *Papio* and were highly correlated with the top PCs when considering baboon samples only (r^2^ > 0.96 between PCs 2-5 including macaques and PCs 1-4 excluding macaques).

To investigate species differences within *Papio*, we subsampled the baboon data to 4 individuals for each species (based on the smallest sample size per species, for Kinda baboons) and analyzed the baboon samples alone. In most subsets (79.6%), PC1 and PC2 mirror the phylogenetic history of the baboon species we sampled (Fig. 1A). They first separated baboons from the northern clade from baboons from the southern clade (PC1), and then separated hamadryas baboons from all other taxa (PC2). To explicitly compare structure in the DNA methylation data to baboon genetic structure, we used Procrustes analyses on the DNA methylation data set and genotype data collected from the same RRBS data (n=49,607 SNPs; Supplementary Methods).

The first two PCs of the genotype data were significantly concordant with the first two PCs of the DNA methylation data (Fig. 1B; Procrustes *t*_0_ = 0.89, p < 10^−6^), indicating that divergence in CpG methylation levels is closely tied to genetic divergence (near-identical results were obtained when including additional PCs, up to PC 6).

Consistent with a close link between DNA methylation and genetic divergence, pairwise genetic covariance between samples strongly predicted pairwise covariance in DNA methylation levels. Across all CpG sites, a sample-wise covariance matrix based on RRBS-derived genotype data was significantly correlated with a sample-wise covariance matrix based on DNA methylation levels (n=756,262 CpG sites; Mantel test r [95% CI] = 0.680 [0.651-0.721], p < 10^−6^), especially when considering the baboon samples alone (r = 0.818 [0.794-0.856], p < 10^−6^).

However, the strength of the correlation varied systematically across genomic contexts (Supplementary Methods). DNA methylation variation among baboons exhibited the lowest correlation with genetic variation in CpG islands (r = 0.497 [0.449-0.554]), gene exons (r =0.594 [0.548-0.688]), and gene promoters (r = 0.602 [0.563-0.652]), and the highest correlation in regions of the genome that are functionally unannotated in *Papio* (r = 0.827 [0.801-0.880]) (all p < 10^−6^). Gene introns, untranslated regions (UTRs), CpG shores, and enhancers fell between these extremes (Fig. 2A). Further, in all contexts, the strongest relationship between genetic variation and DNA methylation levels was observed for intermediately methylated CpG sites, which were also the most variable (Fig. 2A). Notably, regions of the genome that support a non-consensus phylogeny (i.e., those most likely to be affected by incomplete lineage sorting or admixture, which is common in baboons: Zinner, et al. 2009; Zinner, et al. 2013; Tung and Barreiro 2017; Rogers, et al. *in review*; see Methods) exhibited a weaker association between the DNA methylation and genotype matrices than those that fit the consensus phylogeny (Mantel test r = 0.716 [0.649-0.760], n = 211,852 sites compared to 0.815 [0.766-0.858] for regions that matched the consensus phylogeny, n = 542,509 sites).

**Figure 2.**
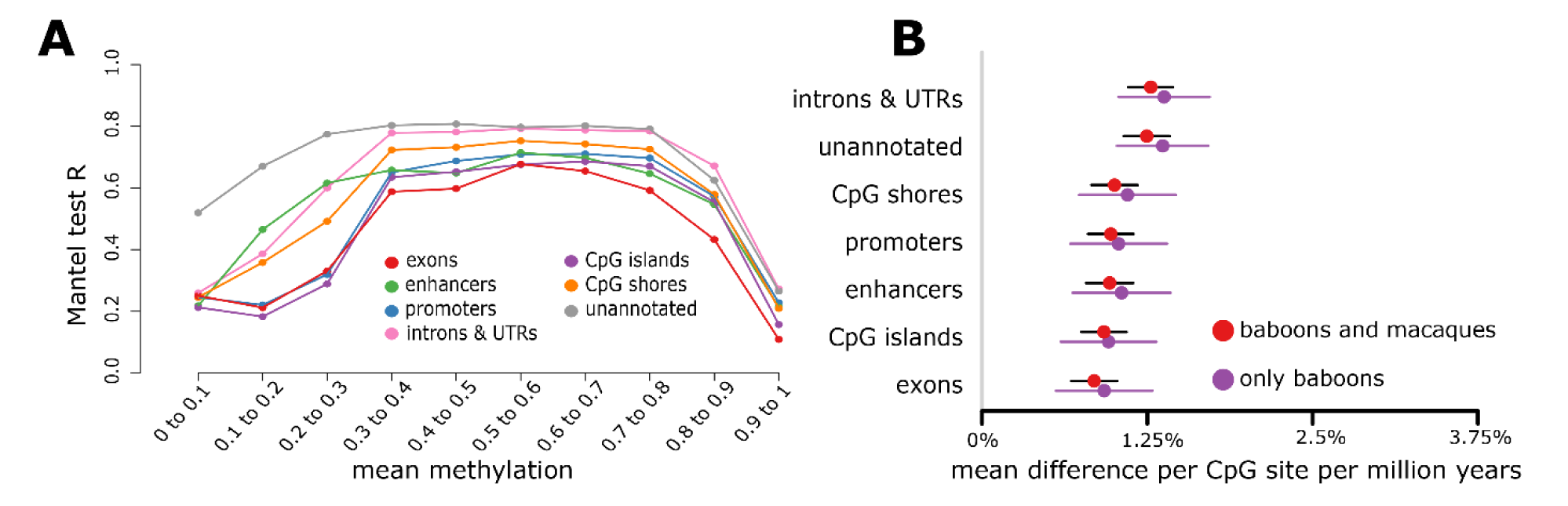
Concordance between DNA methylation variation and genetic variation depends on genomic context. **(A)** Correlation between pairwise genetic covariance between species and pairwise covariance in DNA methylation levels, for CpG sites stratified by genomic context and mean DNA methylation level. Each point represents n=2,658-604,775 CpG sites. **(B)** Estimated mean rate of change in DNA methylation levels per million years, stratified by genomic context. Error bars represent the standard error for each estimate.

Both the PCA results and the correlation between DNA methylation and genetic structure thus suggest that species differences in DNA methylation are largely explained by genetic divergence. To investigate how this relationship scales, we estimated the correlation between divergence time (0.380 – 1.4 million years within *Papio*, and 8.1 million years between baboons and macaques: Perelman, et al. 2011; Rogers, et al. *in review*) and DNA methylation divergence per site. For this analysis, we limited the data set to CpG sites that were measured in at least 3 individuals of each species (n=438,713 CpG sites). When both macaques and baboons were included in the analysis, pairwise divergence time was strongly positively correlated with pairwise DNA methylation divergence (Mantel test r = 0.970, p = 0.011), with an estimated rate of change for the average CpG site of 1.14% per million years. This estimate is similar to that obtained from baboons alone (1.27% per million years), although the baboon results are noisier and not statistically significant (Mantel test r = 0.377, p = 0.067). Divergence in DNA methylation is fastest in functionally unannotated regions of the genome and slowest in gene exons, CpG islands, promoters, and enhancers (Fig. 2B). This pattern is observable whether or not rhesus macaques are included and holds across mean methylation levels, although differences in rate are smaller for sites that are intermediately methylated (Fig. S3).

### Evolutionary shifts in DNA methylation levels within *Papio*

We next investigated the frequency and distribution of CpG sites that exhibit (i) genus-level differences in DNA methylation between baboons and macaques; (ii) clade-level differences in DNA methylation between northern and southern clade baboons; and/or (iii) species-level shifts in DNA methylation levels that differentiate one baboon species from all other baboons. To do so, we first used ANOVA to identify 182,168 (25.2% of those tested), 18,009 (2.5%), and 33,674 (4.7%) CpG sites for which genus, clade (within genus), or species (within clade) membership explained significant variance in DNA methylation levels, respectively (Fig. 3A; 10% FDR: Storey and Tibshirani 2003). These sets of taxonomically structured CpG sites overlapped more than expected by chance (Fisher’s Exact Test log_2_(OR) > 0.75 and p < 10^−16^ for all three pairwise comparisons). CpG sites located in functionally unannotated regions, gene introns, and untranslated regions (UTRs) were more likely to exhibit taxonomically structured variation in DNA methylation than CpG sites in other genomic contexts (Fig. 3B; Table S2). Conversely, such variation was depleted for CpG sites in gene exons. This dependency on genomic context was generally consistent between sites that exhibited significant genus, clade, or species-level variation. However, species-level changes were more strongly enriched in unannotated regions and more clearly depleted for other functional contexts (Fig. S4; Table S3), consistent with faster divergence in regions where genetic variation is more likely to be selectively neutral.

**Figure 3.**
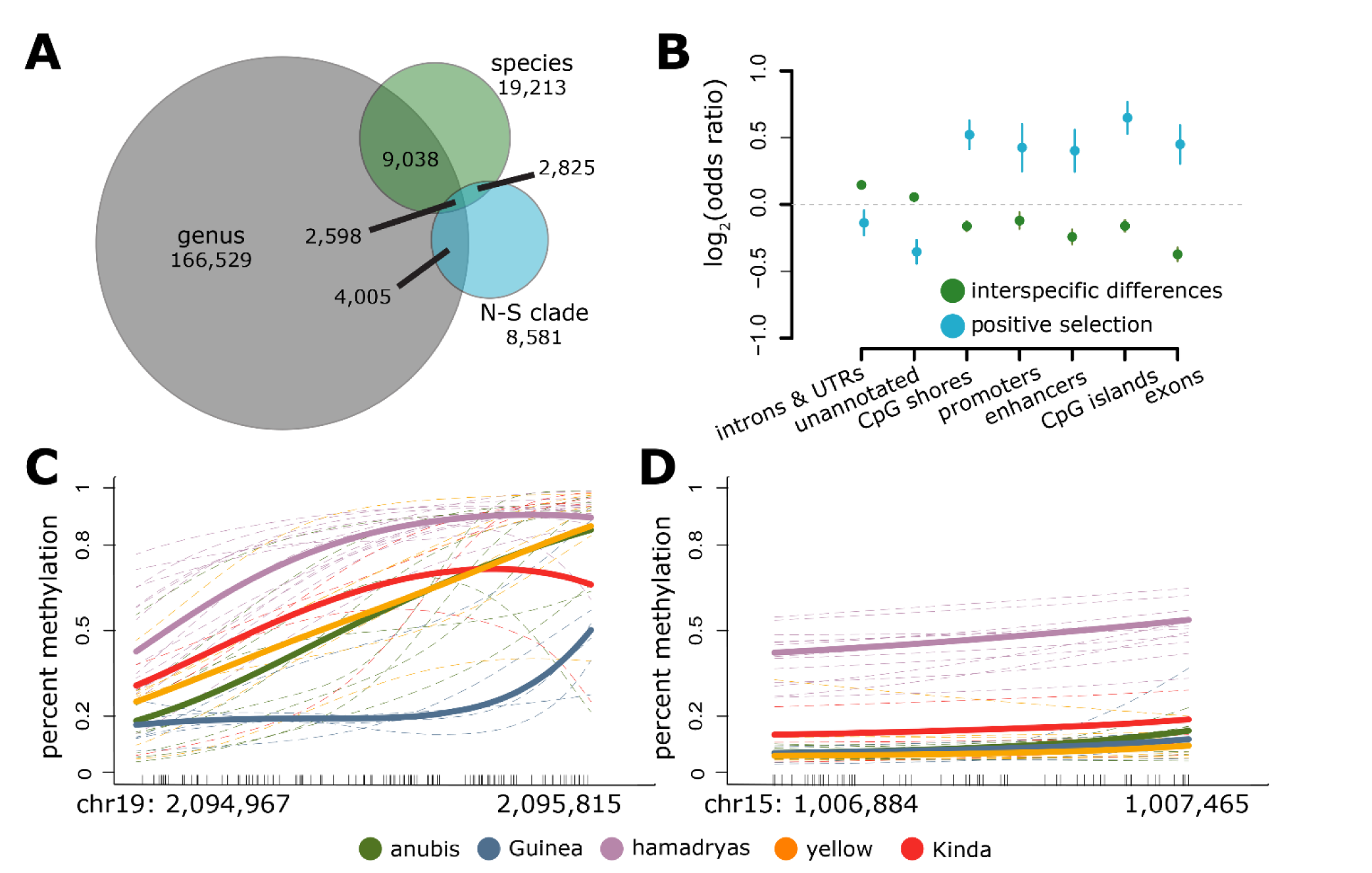
Interspecific differences in DNA methylation levels. **(A)** The number of CpG sites that exhibit significant taxonomic structure at successive levels of the phylogeny. Sites significantly overlap between genus and N-S clade (log_2_(OR) = 0.887, p < 1.28×10^−42^), genus and species (log_2_(OR) = 0.768, p < 2.92×10^−31^), and N-S clade and species (log2(OR) = 3.433, p < 1. 45×10^−295^). **(B)** Enrichment by genomic context for (i) CpG sites in which DNA methylation levels show significant taxonomic structure by clade or species (green dots; background set is the full set of n=756,262 CpG sites analyzed), and (ii) CpG sites in which Ornstein-Uhlenbeck models and heuristic analyses both indicate a likely history of positive selection (blue dots: background set is n=46,260 taxonomically structured CpG sites). Functional elements that are depleted for significant taxonomic structure overall are nevertheless enriched for a signature of selection among those sites that do exhibit taxonomic structure. **(C)** and **(D)** Example large differentially methylated regions (DMRs). Dashes along the x-axis show the location of each measured CpG site in the region and lines show the smoothed mean DNA methylation level (BSmooth: Hansen, et al. 2012). Thin dashed lines represent individual samples, and bold lines represent mean methylation levels per species. A Guinea baboon-specific DMR associated with the *Leucine Rich Repeat and Ig Domain Containing 3* (*LINGO3*) gene is shown in (**C**) and a hamadryas baboon-specific DMR associated with the *taperin* (*TPRN*) and *transmembrane protein 203* (*TMEM203*) genes is shown in (**D**).

To identify shifts in DNA methylation associated with specific baboon taxa, we focused on the set of 46,260 sites that were taxonomically structured by clade or species membership.

For these sites, we then applied a binomial mixed effects model (Lea, et al. 2015) to identify differential methylation (i) between each target species and all other baboons, and (ii) between clades (10% FDR threshold). We required a minimum 10% difference in mean DNA methylation levels between the focal species and all other baboon species to call a species-specific shift, and a minimum 10% difference between all between-clade species pairs, as well as rhesus macaque, to call a clade-level shift. Based on these criteria, we identified 2,959 – 11,189 species-specific shifts per species (29,001 unique sites in all). The number of shifts per species was not a function of sample size or independent evolutionary time (linear model, p = 0.809 and p = 0.743, respectively). We identified another set of 9,803 CpG sites with evidence for a clade-specific shift: 2,843 sites where DNA methylation in the northern clade was different from the southern clade species and macaques, 5,340 sites where DNA methylation in the southern clade was different from the northern clade species and macaques, and 1,640 sites where methylation differed between the two clades and both clades were also different from macaques.

To assess the biological significance of these shifts, we again investigated their distribution across the genome. Relative to the set of 46,260 sites tested, both species- and clade-specific shifts in DNA methylation were depleted in unannotated regions (species: log_2_(OR) = −0.057, p=0.037; clade: log_2_(OR) = −0.139, p = 2.70×10^−5^), suggesting that species- or clade-specific changes are less likely to be neutral than the overall set of taxonomically structured sites. Differentially methylated regions (DMRs, defined as clusters of ≥3 species- or clade-specific sites within a 2 kb window: see Supplementary Methods) were associated with RNA processing and metabolism-related genes in anubis, hamadryas, and yellow baboons, protein targeting in shifts specific to the southern clade, and cell size and organization in shifts specific to the northern clade (10% FDR threshold). We also identified 11 large DMRs (≥ 20 CpG sites: Fig. 3C-D). Six of these DMRs occur in the hamadryas lineage, four in the Guinea lineage, and one is specific to all southern clade baboons. All of the large DMRs overlapped with a CpG island and almost all (9 of 11) were within 10 kb of the nearest gene. Large DMR-associated genes included *single immunoglobulin domain-containing IL1R-related protein* (*SIGIRR*), which is involved in innate immune defense, regulation of inflammation, and natural killer cell maturation; *taperin* (*TPRN*), which is implicated in hearing and sensory phenotypes; and *transmembrane protein 203* (*TMEM203*), which is required for spermatogenesis. These loci represent candidate regions in which differences in DNA methylation may be important in translating genetic variation to phenotypic differences between baboon taxa.

### Selection on DNA methylation patterns in baboons

Our results indicate that DNA methylation in functionally important regions of the genome evolves more slowly than DNA methylation in unannotated regions, consistent with stabilizing selection on gene regulation and neutral evolution for functionally silent CpG sites. However, lineage-specific shifts in DNA methylation also point to a possible contribution of positive selection. To investigate the relative contribution of these different selective regimes, we performed site-specific analyses using two complementary methods: (i) a heuristic approach based on comparisons between intra- and interspecific variation (Rifkin, et al. 2003; Nuzhdin, et al. 2004; Gilad, et al. 2006a; Whitehead and Crawford 2006; Gallego Romero, et al. 2012), and (ii) Ornstein-Uhlenbeck models of phenotypic evolution, which have recently been extended to model gene expression phenotypes and to incorporate intraspecific variation (Lande 1976; Butler and King 2004; Bedford and Hartl 2009; Rohlfs and Nielsen 2014).

The heuristic approach is based on the logic that phenotypes that evolve under positive selection will harbor less intraspecific variation than phenotypes that evolve under genetic drift, (Gallego Romero, et al. 2012). Therefore, CpG sites where mean methylation differs between species but variation is low within species are the most likely to have experienced a history of positive selection. To identify such sites, we focused on those in the lowest decile of within-species variance (controlling for average methylation, see Methods) that also displayed significant species or clade-specific methylation. These criteria yielded a set of 1,178 and 4,399 CpG sites that are candidates for positive selection to differentiate baboon clades or species, respectively. We note that this approach is likely to retain false positives (and also miss false negatives, which is common in tests for selection): thus, this set should be treated as enriched for a likely history of positive selection, rather than as a definitive list of positively selected sites.

In the second approach, we fit Brownian motion and Ornstein-Uhlenbeck models of phenotypic evolution, which include explicit parameters for the strength of selection towards a phenotypic optimum or optima (Butler and King 2004). We used a modified approach that takes into account intraspecific phenotypic variance (following Bedford and Hartl 2009; Rohlfs and Nielsen 2014), with modifications to accommodate our data type. Simulations indicated that, in the baboon phylogeny, these models are underpowered to identify species-specific episodes of selection, but are reasonably well-powered to detect positive selection on multi-species lineages (see Supplementary Methods). However, like the heuristic approach, we treat our results as enriched for specific evolutionary histories, as opposed to definitive. For each taxonomically structured site (n=46,260 sites), we fit five models, which captured (i) genetic drift across the baboon phylogeny; (ii) stabilizing selection towards a single optimum; (iii) positive selection towards a different phenotypic optimum in the southern baboon clade (yellow and Kinda); (iv) positive selection towards a different phenotypic optimum in the northern baboon clade (anubis, Guinea, and hamadryas); and (v) positive selection towards a different phenotypic optimum in a northern baboon subclade, the anubis-Guinea lineage. We defined the best model (for each site) as the one with the lowest Akaike Information Criterion value (AIC: Akaike 1974). Models iii-v, which include positive selection somewhere in the tree, were chosen as the best model for 20,329 CpG sites (2.7% of the initial set of sites tested, n=756,262, and 43.9% of the 46,260 sites that exhibited significant clade- or species-level shifts).

The heuristic approach and the OU model approach produced highly overlapping sets of putative positively selected sites (Fisher’s Exact Test log_2_(OR) = 2.26, p = 8.99×10^−118^). We detected 987 CpG sites with evidence for positive clade-level selection in both methods (Supplementary Methods), which we treat as our highest-confidence set. Compared to the background set of sites with clade- or species-level shifts in DNA methylation, which tend to occur most often in functionally unannotated regions, this set is strongly enriched for gene exons and functional regulatory elements, including promoters, enhancers, CpG islands, and CpG island shores (Fig. 3B; Table S2). This pattern is consistent for all sets of candidate positively selected sites (Fig. S4; Table S4). 129 DMRs were associated with species-specific selection (4-46 per species) and 39 with clade-specific selection (16 assigned to the northern clade and 23 assigned to the southern clade), and consistent with our results for species-specific shifts above, candidate positively selected DMRs were enriched overall for association with genes involved in metabolic processes (10% FDR threshold).

If regulatory divergence in DNA methylation levels is a consequence of genetic divergence, the genetic sequence surrounding positively selected CpG sites should also show signatures of positive selection, including reduced levels of local genetic variation. To test this prediction, we calculated nucleotide diversity (π: Nei and Li 1979) for the 1 kb centered on each taxonomically structured CpG site for each baboon species, based on data from the Baboon Genome Project Diversity Panel (2 – 4 individuals sequenced at 30x coverage per species; see Supplementary Methods). Averaged across all baboon lineages, nucleotide diversity around CpG sites for which we inferred a history of positive selection somewhere in the tree (mean π ± s.d = 0.00239 ± 0.00261) did not differ from nucleotide diversity around CpG sites with no evidence for positive selection (0.00241 ± 0.00221, Tukey’s HSD p = 0.923; Fig. 4). However, nucleotide diversity was significantly lower for site-lineage combinations in which positive selection was specifically inferred than for either other lineages at the same site (0.00213 ± 0.00348 versus 0.00251 ± 0.00273, p = 3.86 × 10^−12^) or near sites with no evidence for positive selection (0.00213 ± 0.00348 versus 0.00241 ± 0.0022, p =1.22 × 10^−12^). For “nonselected” lineages, local nucleotide diversity did not differ from nucleotide diversity at sites with no evidence for positive selection (p = 0.096).

**Figure 4.**
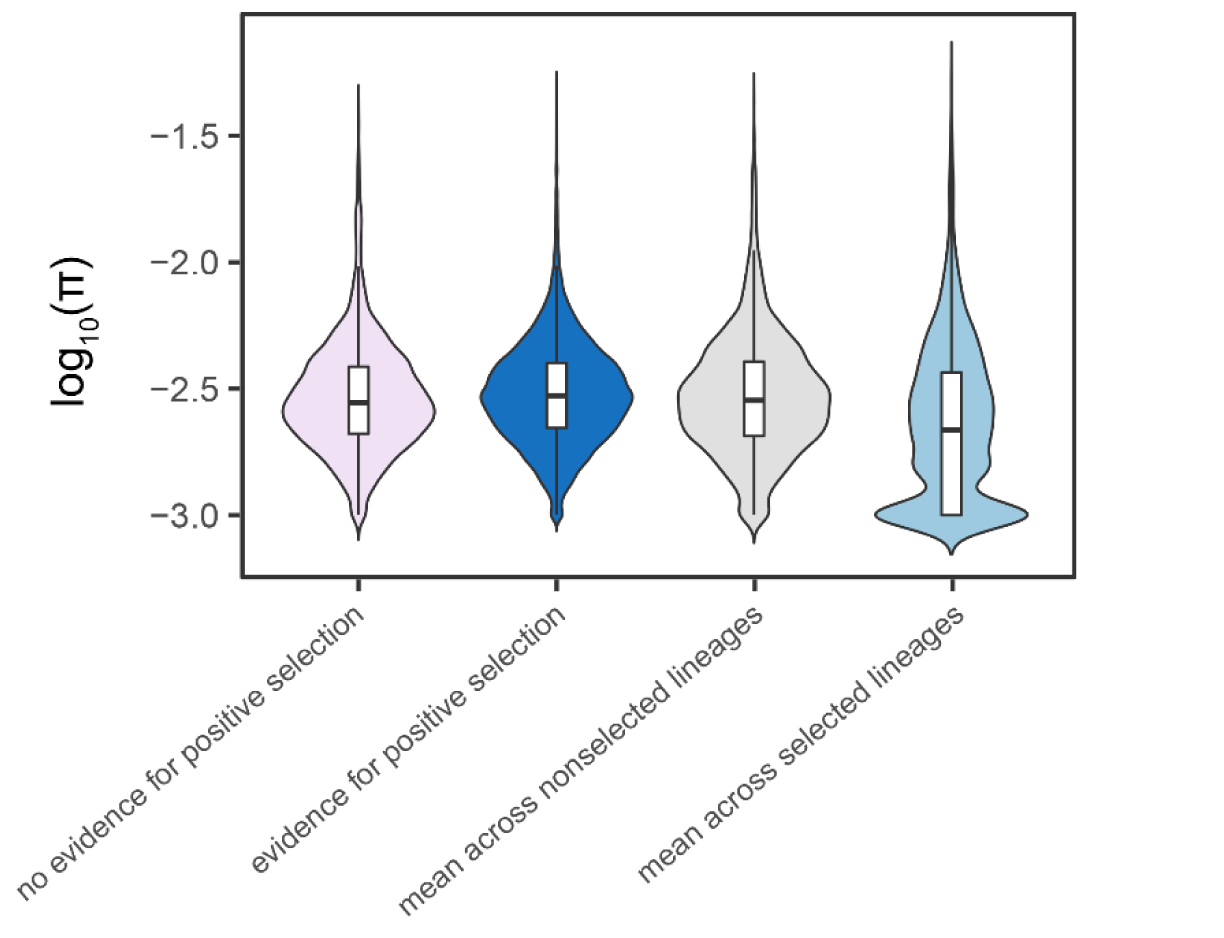
Nucleotide diversity near CpG sites is lower in lineages where positive selection has been inferred. Log_10_(π) for the 1 kb window surrounding CpG sites where DNA methylation levels are taxonomically structured, and (i) we inferred no evidence for positive selection (pink: n=40,212 sites) or (ii) where positive selection was inferred on any baboon lineage (dark blue: n=4,901 sites, based on the intersection set of the heuristic and OU approaches for multi-species lineages and results from the heuristic approach for single lineages). Log_10_(π) for sites in dark blue are replotted in gray for lineages unaffected by putative positive selection and in light blue for lineages putatively affected by positive selection. Lineage-site combinations linked to positive selection (light blue) exhibit lower local nucleotide diversity than all other classes (Tukey’s HSD: p= 1.22 × 10^−12^ compared to sites with no evidence of positive selection [pink]; p = 7.26 × 10^−7^ against the same sites, but with π averaged across all lineages [dark blue]; p = 3.86 × 10^−12^ against the same sites, but with π averaged across nonselected lineages only [gray]). π was calculated separately for each species and averaged across lineages, and is log transformed here for visualization purposes only. Box plots show median (black bar) and interquartile range (whiskers).

## Discussion

Together, our findings provide novel insight into the rate and determinants of DNA methylation divergence in primates. In contrast to comparative studies of human populations (Fraser, et al. 2012; Heyn, et al. 2013; Carja, et al. 2017), but like studies across the more deeply diverged great apes (Hernando-Herraez, et al. 2013; Hernando-Herraez, et al. 2015), global divergence in DNA methylation patterns in baboons is clearly apparent, even among species that diverged relatively recently (e.g., anubis and Guinea baboons: diverged ∼0.38 mya). Roughly speaking, our results suggest that primate taxa can become clearly distinguishable based on DNA methylation data after approximately 35,000 generations (assuming a generation time for baboons of 11 years: Swedell 2011; Rogers, et al. *in review*), although this rate varies by genomic context. Notably, although yellow baboons and Kinda baboons diverged earlier than anubis and Guinea baboons (∼0.6 mya, closer to when hamadryas baboons diverged from the anubis-Guinea lineage), global patterns of DNA methylation separate these two southern clade species less clearly than any of the northern clade species. This difference may reflect recent admixture in the southern part of the yellow baboon range (Zinner, et al. 2009; Keller, et al. 2010), or smaller long term effective population sizes in the northern clade species (Rogers, et al. *in review*). Among the northern clade species, anubis baboons fall closest to southern clade baboons, which may also be a consequence of hybridization: anubis baboons and yellow baboons hybridize in Kenya today, and have likely done so in the past as well (Alberts and Altmann 2001; Charpentier, et al. 2012; Wall, et al. 2016; Rogers, et al. *in review*).

Our results are in line with emerging evidence that, in comparisons involving clearly divergent lineages, variation in DNA methylation levels is largely tied to variation in nearby genetic sequence (Hernando-Herraez, et al. 2013; Hernando-Herraez, et al. 2015). Specifically, DNA methylation patterns in baboons recapitulate phylogenetic structure, and local genomic context predicts both the rate at which DNA methylation evolves and the probability of a past history of selection. These observations are consistent with analyses in great apes, which revealed that interspecific differences in DNA methylation tend to occur at loci that also contain high levels of species-specific mutations (Hernando-Herraez, et al. 2015). Similarly, in *Arabidopsis* lines, inter-accession differences can largely be explained by *cis*-acting methylation quantitative trait loci (meQTL) (Dubin, et al. 2015). Thus, while environmental variation may be important for explaining variation in DNA methylation within populations (Jirtle and Skinner 2007; Feil and Fraga 2012), including baboons (Lea, et al. 2016), genetic effects are likely to dominate in between-population and between-species comparisons. Indeed, in our data set, hamadryas baboon and anubis baboon samples were obtained from multiple populations, representing both captive and natural settings. However, despite exposure to different diets and housing conditions, population differences explained very little variance in the overall data set (Supplementary Methods).

Our data set also facilitates initial comparisons of DNA methylation evolution against gene expression data sets. Although our findings resemble those of cross-species gene expression analyses in that they globally reproduce the species phylogeny, they also suggest that the evolution of DNA methylation is less constrained on average. While CpG sites are enriched in gene bodies, promoters, and CpG islands, the majority of CpG sites in primate genomes fall in functionally unannotated regions. Our analyses show that DNA methylation levels in unannotated regions are both faster evolving, and, compared to all rapidly evolving sites, underrepresented for signatures of positive selection (Fig. 3B). Thus, while several lines of evidence indicate that gene expression levels for most genes are constrained by stabilizing selection, the same pattern probably does not hold for most CpG sites. This difference may explain why the evolution of DNA methylation levels looks more clock-like than for gene expression (Carja, et al. 2017), a pattern now observed in human populations, *Arabidopsis* accessions, and here, in baboons (Becker, et al. 2011; Schmitz, et al. 2011; van der Graaf, et al. 2015; Carja, et al. 2017). It also is consistent with experimental studies showing that DNA methylation levels influence gene regulation at only a subset of CpG sites (Maeder, et al. 2013; Ford, et al. 2017; Lea, et al. 2017b).

Nevertheless, we do find support for positive selection on DNA methylation levels for a small fraction of the CpG sites we profiled. Tests for selection on phenotypic variation have important limitations (e.g., unknown mutational variance, the assumption of relatively simple evolutionary scenarios: Butler and King 2004; Gilad, et al. 2006b; Rohlfs and Nielsen 2014).

However, they are still likely to enrich for true cases of positive selection (Blekhman, et al. 2008; Rohlfs and Nielsen 2014). Here, the strong enrichment of putatively selected sites within genes and gene regulatory elements, the overlap between two different methods for identifying selected sites, and the identification of coherent DMRs associated with candidate selected sites all indicate that we have captured a set of CpG sites of interest for baboon evolutionary history.

Additionally, we identified a loss of local nucleotide diversity—a purely DNA sequence-based analysis—specifically near sites and in lineages inferred to be affected by positive selection, in an analysis based only on DNA methylation phenotypes.

Recent evidence shows that changes in DNA methylation can play an important role in phenotypic evolution. For example, loss of sight in cave-dwelling tetra fish (*Astyanax mexicanus*) is mediated by DNA methylation-mediated repression of genes involved in eye development (Gore, et al. 2018). Our results suggest that comparative studies of DNA methylation in recent radiations can help identify other loci of interest, and could potentially be combined with outlier scans based on other types of data (e.g., Bergey, et al. 2016). Notably, in baboons, we found several large DMRs linked to genes involved in immunity, sensory perception, and spermatogenesis, three categories previously identified in sequence-based scans for selection in primates (Kosiol, et al. 2008). These examples suggest that, at least in some instances, natural selection on gene regulation has been directed towards changes in DNA methylation phenotypes. If so, variation in DNA methylation at candidate selected sites should functionally affect gene expression, a prediction that can now be empirically tested using reporter assays or epigenomic editing approaches (Liu, et al. 2016; Lea, et al. 2017b). We anticipate that such a combination of comparative, genetic, and experimental approaches will help resolve the much-debated role of epigenetic marks in adaptive evolution (Laland, et al. 2014; Verhoeven, et al. 2016).

## Methods

### RRBS data generation, processing, and quality control

DNA methylation data were generated for 39 baboons across five of the six recognized extant species (9 anubis, 6 yellow, 14 hamadryas, 6 Guinea, and 4 Kinda baboons; Table S1). We also generated RRBS data for 5 rhesus macaques as an outgroup. For *P. anubis* samples from the Washington National Primate Research Center (WaNPRC), *P. papio* from the Brookfield Zoo, and *P. hamadryas* from the North Carolina Zoo, we extracted genomic DNA using the QIAGEN DNeasy Blood & Tissue Kit, following the manufacturer’s recommendations. Other samples were obtained as previously extracted DNA (see Table S1). All DNA samples were extracted from whole blood with the exception of 2 *P. cynocephalus*, 1 *P. anubis*, 2 *P. kindae*, and 1 *P. hamadryas* for whom samples were obtained from banked white blood cells.

Differences in source tissue (whole blood versus banked white blood cells) do not contribute to any of the first 10 principal components of variation in DNA methylation within this sample (t-test, all p-values > 0.20). Differences in cell type composition also appear unlikely to drive species-specific methylation levels (Supplementary Methods).

RRBS libraries for each sample were prepared following Boyle et al. (2012). Briefly, Illumina TruSeq barcoded libraries were constructed using 180 ng of genomic DNA per sample. Libraries were pooled together in sets of 10-12 samples, subjected to sodium bisulfite conversion using the EpiTect Bisulfite Conversion kit (QIAGEN), and then PCR amplified for 16 cycles prior to sequencing on the Illumina HiSeq 2500 platform. Each pooled set of libraries was sequenced in a single lane to 17.2 million reads per sample (s.d. = 12.8 million reads: Table S1). To assess the efficiency of the bisulfite conversion, 1 ng of unmethylated lambda phage DNA (Sigma Aldrich) was added to each sample prior to library construction.

Sequences were trimmed for adapter contamination, RRBS end repair, and base quality using Trim Galore! (Babraham Bioinformatics) before being mapped to the anubis baboon reference genome (*Panu2.0*) using BSMAP (Xi and Li 2009). We removed sites that overlapped genetic variants in which one allele abolishes a CpG site found in the reference genome.

Combined with BSMAP’s three-nucleotide mapping option, this step eliminates most heterospecific mapping biases within *Papio* (Supplementary Methods and Fig. S5). The DNA methylation level at each CpG site was calculated as the proportion of reads with unconverted (i.e. methylated) cytosine bases to total reads covering that site. Based on reads mapped to the lambda phage genome, all samples had a bisulfite conversion efficiency greater than 98.5%, with no significant contribution of species identity to variance in conversion efficiency (ANOVA F = 1.303, p = 0.27; Table S1).

After excluding sites for which data were missing for ≥50% of our study subjects or for which mean coverage was <5x, we retained 2,450,153 CpG sites for downstream analysis. As expected for RRBS data sets, these sites were enriched in functionally important regions of the genome and displayed typical mammalian patterns of CpG DNA methylation (Fig. S1). To focus on the sites most likely to exhibit biologically meaningful variation, we further excluded constitutively hypermethylated (mean DNA methylation level >0.90) and constitutively hypomethylated (mean DNA methylation level <0.10) sites and those that were near-invariant (s.d. < 0.05), resulting in a final analysis set of 756,262 CpG sites.

Where possible, we modeled DNA methylation levels as count data (the number of methylated reads and total reads for each site), which retains information about the uncertainty in each estimate due to variation in read coverage (Dolzhenko and Smith 2014; Sun, et al. 2014; Lea, et al. 2015; Lea, et al. 2017a). However, because some of our analyses (e.g., PCA, Ornstein-Uhlenbeck models) required continuous data, we also estimated DNA methylation levels as the ratio of methylated reads to total reads within each individual for each CpG site. Because variation in sequencing coverage can systematically bias DNA methylation estimates, for these analyses we used the residuals of the raw ratios after regressing out site-specific total read coverage for each individual.

### Functional element annotations and enrichment analysis

We used gene body and CpG island annotations for *Panu2.0* obtained from Ensembl (Cunningham et al. 2015) and the UCSC Genome Browser (Karolchik et al. 2014), respectively. Gene promoters were defined as the 2 kb region upstream of the 5’-most annotated gene transcription start site (following Deng et al. 2009; Shulha et al. 2013; Lea et al. 2015) and CpG island shores were defined as the 2 kb regions flanking either side of a CpG island (Irizarry, et al. 2009). Because baboon enhancer annotations are not available, we defined putative baboon enhancers by projecting coordinates from ENCODE H3K4me1 ChIP-seq of human peripheral blood mononuclear cells (Dunham et al. 2012) onto the *Panu2.0* genome using the UCSC Genome Browser *liftover* tool (Hinrichs et al. 2006).

Geneontology (GO) enrichment analyses were performed using the Cytoscape module ClueGO (Bindea, et al. 2009). To link differentially methylated sites to genes, we first identified clusters of CpG sites with similar patterns of differential methylation (differentially methylated regions or DMRs). We called DMRs when ≥3 CpG sites within a 2 kb window exhibited the same type of lineage-specific change (e.g., hypo-methylation in hamadryas baboons), and bounded the DMR by the first and last CpG site that exhibited lineage-specific methylation. We then collapsed overlapping DMRs. We assigned a DMR to a gene when a CpG site within the DMR fell within 10 kb of the gene body. To test for gene set enrichment, we analyzed GO Biological Processes that fell between levels 3 and 8 of the GO tree, included at least 4 genes in our data set, and for which at least 5% of genes assigned to the term were present in the test set. We also collapsed GO parent-child terms with at least 50% overlap. Enrichment analyses were corrected for multiple hypothesis testing using the Benjamini-Hochberg (B-H) method (Benjamini and Hochberg 1995). Gene set enrichment analyses for DNA methylation data can be biased if some gene sets are systematically associated with larger numbers of CpG sites than others (Geeleher, et al. 2013). However, in our data set, genes associated with differentially methylated sites were not associated with more tested sites than other genes (logistic regression: z = 0.054, p = 0.957).

### Covariance between genetic structure and DNA methylation patterns

To assess the relationship between phylogenetic structure and DNA methylation patterns in our data set, we conducted principal components analysis in R (version 3.2.5; R Core Team 2016) on the scaled variance-covariance matrix of the DNA methylation level data. We ran the PCA both including and excluding the rhesus macaque samples, and in baboons after subsampling to the same number of individuals per species (n=4; Fig. 1A, 1B, and S2).

To test the correlation between DNA methylation levels and pairwise genetic distance between samples, we used Mantel tests. We called genotypes from RRBS data for 49,607 biallelic SNPs (see Supplementary Methods) and calculated the pairwise genetic covariance. We then compared a genotype-based covariance matrix to the pairwise covariance of DNA methylation profiles using the R package *vegan* (Oksanen et al. 2016), stratified by both functional compartment (gene, enhancer, CpG island, CpG shore, promoter, unannotated) and mean methylation level (Fig 2A). We also tested whether windows of the genome where genetic structure followed an alternate phylogeny (a consequence of incomplete lineage sorting or admixture) exhibited a lower correlation between genetic and DNA methylation covariance (see Supplementary Methods).

Finally, to investigate the relationship between DNA methylation divergence and genetic divergence between species, we retained CpG sites for which each species was represented by at least three individuals and a total (across individuals) of at least 10 reads (n = 438,713 CpG sites). We calculated the mean DNA methylation level per species for each retained CpG site and the difference in mean methylation between each species pair. We then tested whether divergence time (based on Rogers, et al. *in review* for baboons and Perelman, et al. 2011 for baboon-macaque) predicted the Euclidean distance between species using a Mantel test.

### Lineage-specific changes in DNA methylation

For sites in which clade or species significantly contributed to variance in DNA methylation levels (n=46,260 taxonomically structured sites, identified using ANOVA and a 10% FDR threshold), we tested for lineage-specific shifts using the beta-binomial model implemented in the program MACAU (Lea, et al. 2015). We tested each species for differences in DNA methylation level when compared to all other baboons and we also tested whether southern clade baboons had different methylation levels than northern clade baboons. For each comparison and CpG site, we considered the model:

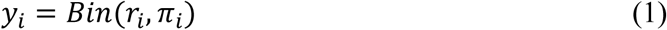

where *r*_*i*_ is the total read count for *i*^th^ individual, *y*_*i*_ is the methylated read count for that individual, and *π*_*i*_ is an unknown parameter that represents the true methylation level for that individual at the site of interest. MACAU then uses a logit link to model *π*_*i*_ as a function of the predictor variable of interest (here, species or clade membership):

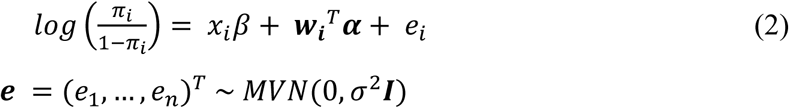

where ***w_i_*** is a vector of fixed effect covariates including an intercept and the sample-specific bisulfite conversation rate; *α* is a vector of coefficients for ***w*_*i*_**; *x*_*i*_ represents species or clade membership coded as 1 (for the taxon of interest) or 0 (for any other taxa) and *β* is the coefficient for the effect of taxonomic membership; ***e*** is an *n*-vector of independent residual error with variance *σ*^2^; and ***I*** is a n-by-n identity matrix. We did not model genetic non-independence in this analysis; thus, the ***K*** matrix input to MACAU was an identity matrix.

In addition to a 10% FDR threshold (q-value: Storey & Tibshirani 2003), we required a minimum difference of 10% in mean methylation between either (i) the focal species compared to all other species, for species-level shifts, or (ii) for all pairwise comparisons between northern clade and southern clade species, for clade-level shifts. We assigned clade-level shifts to one of the two lineages based on post-hoc comparison to rhesus macaques. For example, we assigned a shift to the northern clade when there was a mean difference in DNA methylation of ≥10% between northern clade baboons and macaques, but not between southern clade baboons and macaques.

### Identification of candidate directionally selected sites

To test for positive selection using the heuristic approach, we first calculated the intraspecific variance for each of the 756,262 CpG sites in our primary data set, after mean-centering DNA methylation levels for each species. We then binned the CpG sites into 5% quantiles based on mean methylation level, and retained sites with intra-specific variance in the lowest 10% quantile for each bin. We intersected these low-variance sites with the set of sites that exhibited species- or clade-specific methylation, based on the criteria outlined for identifying taxonomic structure with ANOVA followed by beta-binomial regression. This intersection set is likely to be enriched for a history of positive selection.

As an alternative approach, we fit Ornstein-Uhlenbeck (OU) models of the evolutionary process, based on the phylogenetic tree for baboons (Rogers, et al. *in review*). In OU models, trait evolution is modeled as the sum of stochastic and deterministic forces, with parameters for the strength of selection, the strength of genetic drift, and the trait optimum. In addition, because these models assume phenotypes have a continuous distribution, we transformed DNA methylation levels using a logit link function. A basic OU model has the form:

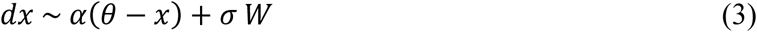

where *dx* captures the continuous rate of change in the trait value *x*, *α* represents the pull towards the optimum trait value *θ*, *σ* is the rate of neutral drift, and *W* is distributed normally with variance corresponding to the amount of independent evolutionary time, *dt*. For multiple species *m*, the OU process can be written as a multivariate normal distribution:

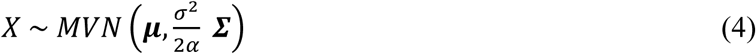

where ***μ*** is a *m*-by-1 vector of *θ*_*j*_, the optimum trait values for species *j*. Σ captures the covariance between species and is determined by the phylogenetic covariance, ***Σ_phylo_***, and *α* such that the covariance between species *j* and *k*, Σ_*j,k*_, is given by 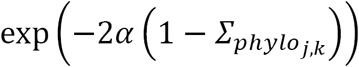. To incorporate intraspecific variance into the OU process, which increases the power to identify true instances of positive selection (Rohlfs and Nielsen 2014), the vector ***μ*** is expanded to an *n*-by-1 vector where each element, *θ*_*i*_, is equal to *θ*_*j*_ for the species *j* to which individual *i* belongs. The covariance matrix ***Σ*** is replaced by the *n*-by-*n* covariance matrix between individuals, with a new parameter *τ*^2^ added to the diagonal of the covariance matrix to take into account within-species variance.

Different evolutionary regimes correspond to different OU process parameter values. Values of *α* at or near 0 correspond to genetic drift (no pull towards an optimum trait value), while non-zero values of *α* indicate a history of selection. If *α* > 0 and *θ* is constant across lineages, the trait has evolved under stabilizing selection. If *α* > 0 and *θ* varies between lineages, the trait has evolved under directional (positive) selection on at least part of the phylogenetic tree. We therefore used AIC to compare five OU models for each CpG site in which species or clade membership significantly contributed to DNA methylation variation based on ANOVA (see Results: *Selection on DNA methylation patterns in baboons* and Supplementary Methods for simulation results on power to detect selective shifts).

## Acknowledgements

We gratefully acknowledge Justin O’Riain, Laurel Serieys, Christian Roos, Julia Fischer, Dietmar Zinner, Jane Phillips-Conroy, Mark Wilson, and the North Carolina Zoo for contributing samples. We thank Amanda Lea for her assistance in generating and analyzing the DNA methylation data used in this study, Rori Rohlfs for valuable aid in using OU models, and members of the Tung lab for helpful comments and discussion. The work reported in this paper also benefited from the activities and discussions within the Baboon Genome Analysis Consortium, for which the full list of participants is presented in Rogers, et al. (*in review*).

Finally, we thank the Baylor College of Medicine Human Genome Sequencing Center for access to the baboon genome assembly (*Panu* 2.0). RRBS data are deposited in NCBI’s Short Read Archive (GSE 4359166). Code for fitting OU models is available at https://github.com/TaurVil/PapioMethylation. This work was supported by the National Science Foundation (grant number BCS-1751783) to J.T. and T.P.V., the National Institutes of Health (R01-GM102526 to J.T. and P51-OD011132 in support of Yerkes National Primate Research Center), a pilot award from the National Center for Advancing Translational Sciences (grant number UL1TR001117), and high-performance computing resources supported by the North Carolina Biotechnology Center (grant number 2016-IDG-1013).

